# Surveillance of CKD epidemiology in the US – a joint analysis of NHANES and KEEP

**DOI:** 10.1101/292698

**Authors:** OB Myers, VS Pankratz, KC Norris, JA Vassalotti, M Unruh, C Argyropoulos

## Abstract

Chronic Kidney Disease (CKD), is highly prevalent in the United States. Epidemiological systems for surveillance of CKD rely on data that are based solely on the NHANES survey, which does not include many patients with the most severe and less frequent forms of CKD. We investigated the feasibility of estimating CKD prevalence from the large-scale community disease detection Kidney Early Evaluation and Program (KEEP, *n* = 127,149). We adopted methodologies from the field of web surveys to address the self-selection bias inherent in KEEP. Primary outcomes studied were CKD Stage 3-5 (estimated glomerular filtration rate [eGFR] <60 mL/min/1.73m^2^, and CKD Stage 4-5 (eGFR <30 mL/min/1.73m^2^). The unweighted prevalence of Stage 4-5 CKD was higher in KEEP (1.00%, 95%CI: 0.94-1.05%) than in NHANES (0.51%, 95% CI: 0.43-0.59%). Application of a selection model with IPW of variables related to demographics, recruitment and socio-economic factors resulted in estimates similar to NHANES (0.55%, 95% CI: 0.50-0.60%). Weighted prevalence of Stages 3-5 CKD in KEEP was 6.45% (95% CI: 5.70 7.28%) compared to 6.73% (95% CI: 6.30-7.19%) for NHANES. Application of methodologies that address the self-selection bias in the KEEP program may allow the use of this large, geographically diverse dataset for CKD surveillance.

## INTRODUCTION

Chronic Kidney Disease (CKD) is a growing public health concern in the United States due to its high prevalence (~13% of the adult US population), association with increased morbidity, mortality and progression to End Stage Renal Disease (ESRD). Treating CKD and ESRD and their complications is extremely costly, accounting for more than 20% of fee-for-service Medicare spending and over 80 billion dollars in the US for 2013 alone (1). Epidemiological systems for geo-temporal CKD surveillance could both fulfil public health objectives (2,3), and direct research efforts towards a better understanding of localized “hotspots” of CKD similar to what has been reported in other contexts (4).

Efforts to develop such a project (2,3) culminated in the establishment of the Center for Diseases Control surveillance project (5), which provides data on CKD incidence, prevalence, disease awareness and other disease indicators. Despite the wealth of data incorporated in the CDC project, prevalence estimates in the general population are based solely on the NHANES survey(5–9). An important limitation of this feature is that individuals with the most severe and costly, but less frequent stages of CKD (4–5) are not well represented in NHANES (7).

Incorporating additional, larger data sources that are more likely to ascertain persons with more severe forms of CKD has the potential to enhance our existing CKD surveillance infrastructure. In this report, we examine the feasibility of estimating CKD prevalence using data from the large community disease detection program of the National Kidney Foundation’s Kidney Early Evaluation and Program (KEEP)(10) in juxtaposition to the population-based data from NHANES.

As a voluntary detection program, KEEP is likely to suffer from a substantial degree of self-selection bias. If it is possible to address this bias, KEEP would provide a population-level data source on all aspects of CKD, including the less common but costly advanced stages of the disease. Use of a national reference population to standardize a sample is used routinely in public health surveillance(11,12). Fueled by the expansion of internet based data collection strategies, there has been a growing literature regarding the handling of self-selection effects in voluntary surveys (13,14) by matching against a reference, representative, probability sample. To our knowledge, similar techniques have not been applied to kidney disease research. In this report, we address the self-selection bias in the KEEP dataset by developing selection models based on subject level factors related to recruitment (selection) probabilities in KEEP. We accomplish this by estimating propensities for KEEP participation relative to NHANES, and using them to form inverse-probability weights (IPW), the application of which adjust the observed percentages of CKD prevalence from this self-selected sample. We demonstrate that this approach can make the estimated CKD prevalence rate from KEEP directly comparable to those of NHANES, opening up the possibility of using this large, geographically diverse dataset for the purpose of CKD surveillance.

## RESULTS

### Baseline characteristics and selection effects in KEEP

The KEEP study made more than 185,000 assessments from August 2000 to June 2013. We analyzed the initial encounter for program eligible KEEP participants assessed by 48 regional affiliates from 2001 to the end of 2012 using data provided by NKF. Participants were excluded if <20 years of age, on dialysis, were pregnant, had a previous kidney transplant, were lacking a valid state of residence or had other invalid demographic variable values (32,026 records excluded, Figure 1). Another 6,317 records that lacked a valid eGFR stage determination were excluded to obtain 127,876 participants (147,168 records) with CKD stage. We then removed follow up visits (19,292 records) and participants seen before 2001 (n = 727) to obtain a sample of 127,149 KEEP participants for propensity score estimation. Our original NHANES sample from 2001 – 2012 had 59,423 participants attending an MEC. We excluded NHANES participants <20 years old (*n* = 27,796), pregnant women (*n* = 963), hemodialysis patients (*n* = 114) and participants without a valid eGFR determination (n = 2,033) to obtain a sample with 28,517 NHANES participants for propensity score estimation.

**Figure 1.**
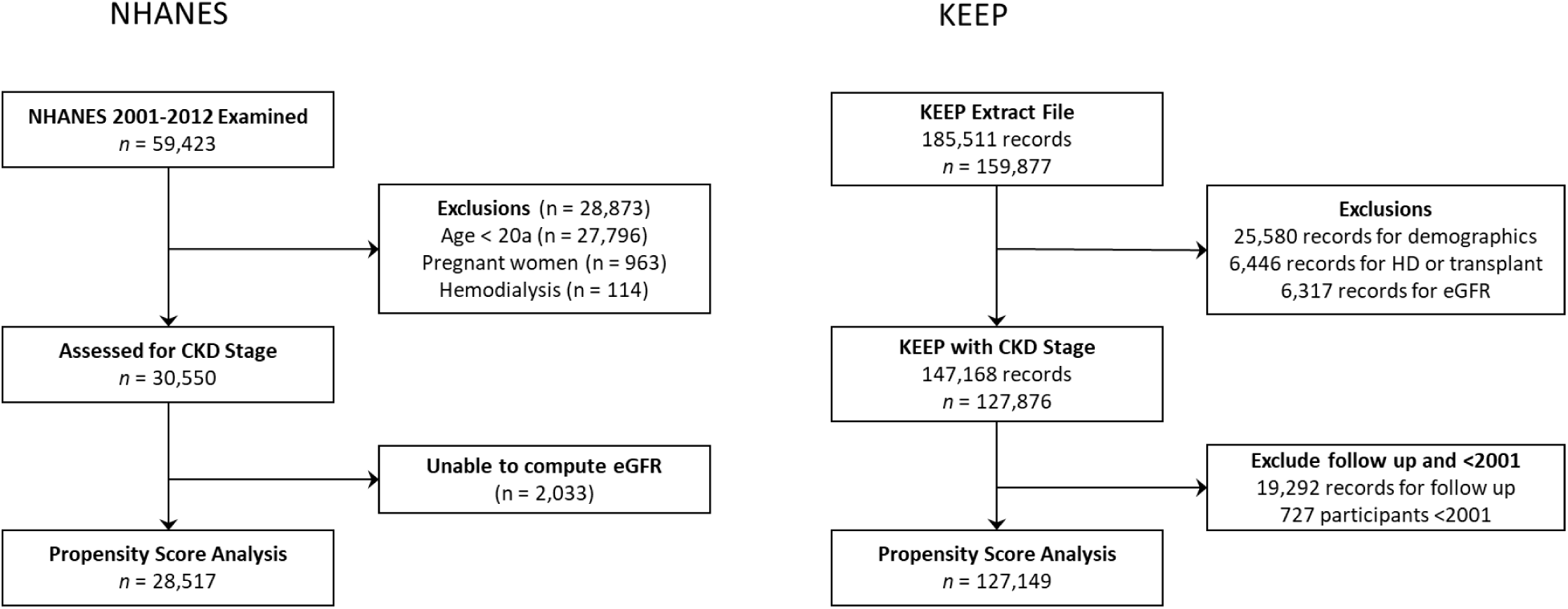
National Health and Nutrition Examination Survey (NHANES) and Kidney Early Evaluation Program (KEEP) participation in study.

KEEP participant characteristics are compared to NHANES general population estimates in Table 1. KEEP participants were older, more likely to be female, less likely to be non-Hispanic White but more likely to be Black than NHANES. KEEP recruitment resulted in an oversampling relative to NHANES of participants with self-reported diabetes, hypertension, CKD, CHF/CAD/Stroke, a family history of diabetes and of heart attack. KEEP participants were also more likely to have reported being obese, to be uninsured and to have at least a high school education. KEEP participants were less likely to be current smokers than the general population. The boosted CART model that produced the best agreement between expected and predicted KEEP frequencies was the model with four-way interactions. Prevalence of CKD within the KEEP sample by IPW decile shows the success of KEEP affiliates recruiting participants at a higher risk of CKD. Smaller deciles (higher probability of selection into KEEP) have significantly higher prevalence of CKD Stages 3-5 than large deciles (*P* < 0.001, Supplemental Figure 2). After KEEP summaries were estimated using weights from this model, frequencies were much closer to NHANES (Table 1). Before weighting the youngest and oldest age categories were under and over represented in KEEP by -19% and 16.6%, but after weighted estimations all categories were within 1.1% of NHANES. Females were 51.1% of NHANES and 52.2% in weighted KEEP compared to 67.7% of the unweighted KEEP sample. Weighted KEEP estimates were also much closer to NHANES for race/ethnicity categories so that all groups were within 2% of expected. Differences between weighted KEEP percentages and NHANES that were more than ±1% included fewer self-reported diabetes (-1.1%), more hypertension (1.9%), family history of diabetes (3.2%), overweight (2.7%), obese (2.9%) and insured (1.4%).

**Table 1.**
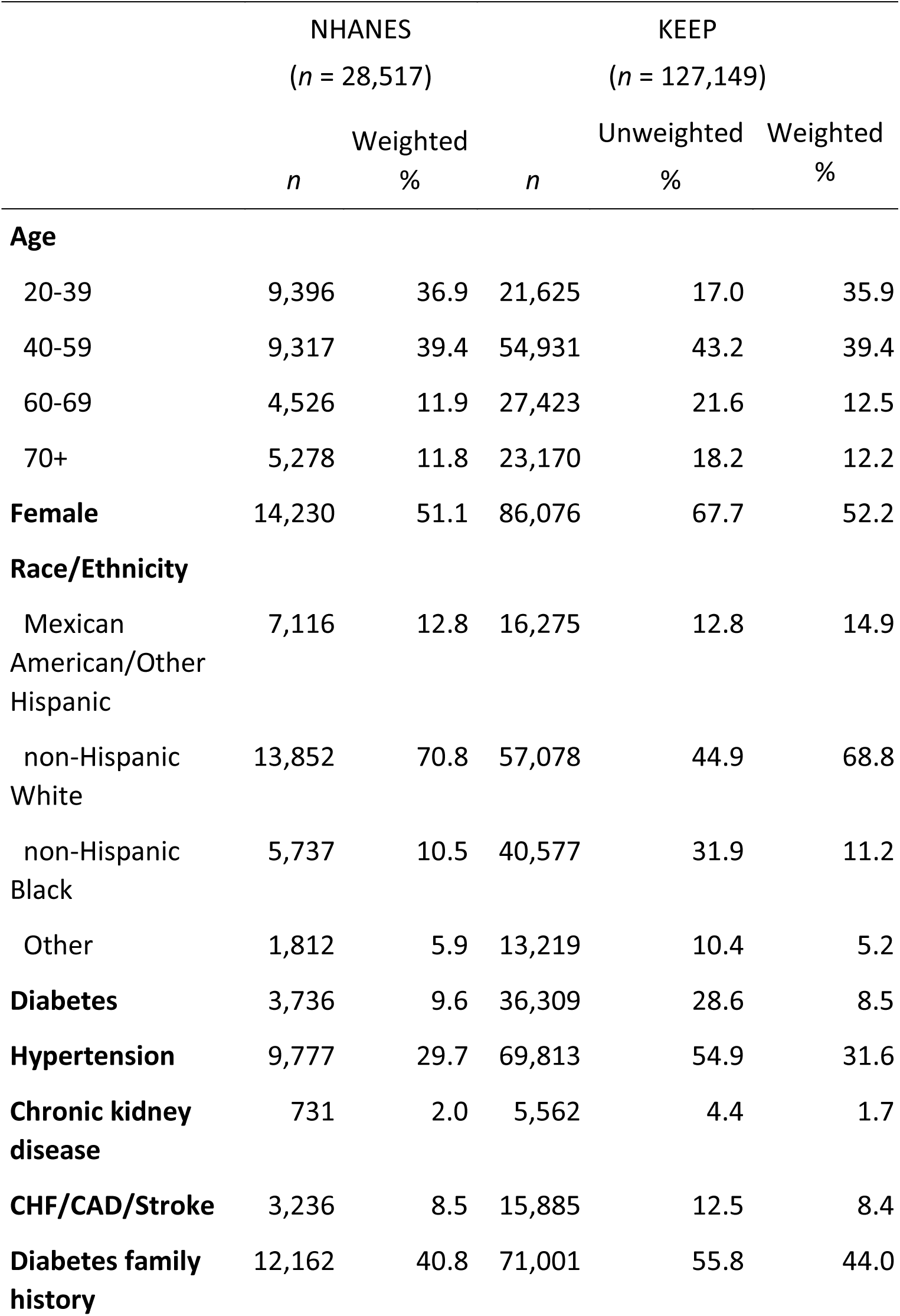
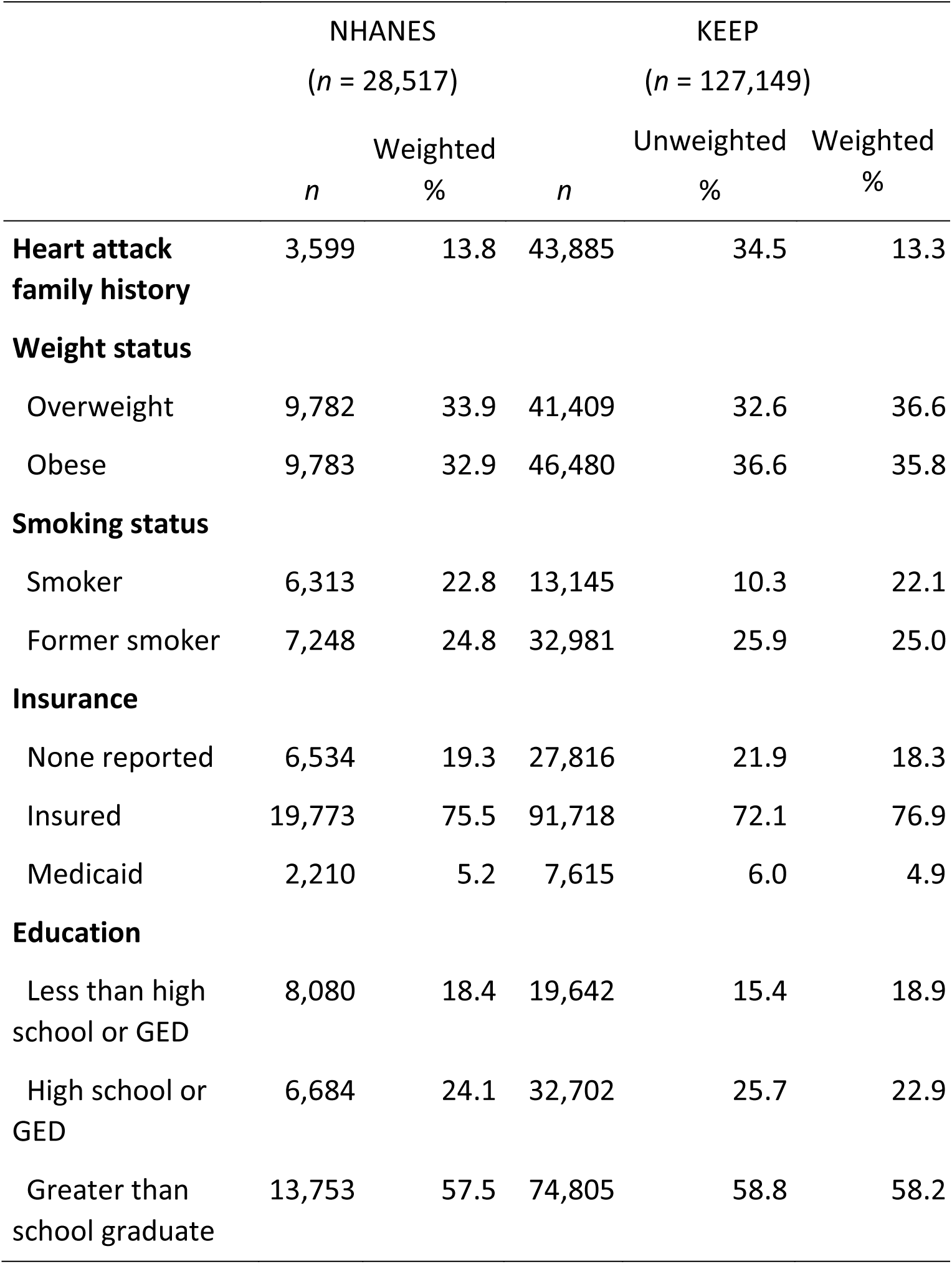
Self-reported characteristics of NHANES reference population and KEEP participants without and with inverse probability weighting, 2001-2012. CHF/CAD/strke - congestive heart failure/coronary artery disease or stroke; CKD - chronic kidney disease; KEEP - Kidney Early Evaluation Program; NHANES - National Health and Nutrition Examination Survey

### Prevalence of CKD stages in NHANES and KEEP

Prevalence of CKD stage 3 in the general population age 20 and older for 2001-2012 was 6.23% (95% CI: 5.82-6.66) from NHANES (*n* = 2,453 cases) compared to 13.14% (95% CI 12.95-13.33) in the KEEP sample during the same period (Table 2, *n* = 16,706 cases). Simple reweighting the KEEP data reduced the prevalence estimate by more than one third to 8.57% and accounting for clustering of KEEP samples within regional affiliates reduced the estimate to 6.45% (5.70-7.28%, Table 2 Weighted-GEE). The average prevalence of CKD Stage 4-5 among all participants was 0.51% in the NHANES general population (95% CI 0.43-0.59%, *n* = 239 cases) and was 1.00% in the KEEP sample (95% CI 0.94-1.05%, Table 2, *n* = 1,267 cases). IPW adjustment accounting for affiliate clusters removed almost all bias so the weighted KEEP prevalence was 0.52% (95% CI 0.42-0.64%, Weighted-GEE).

**Table 2.** CKD prevalence estimated from NHANES and from KEEP without and with inverse probability weighting, 2001 - 2012. Weighted estimates accounted for NHANES sampling design or self-selection by KEEP. KEEP GEE estimates accounted for participant clustering within regional affiliates. CKD - chronic kidney disease; KEEP - Kidney Early Evaluation Program; NHANES - National Health and Nutrition Examination Survey

|  |  | NHANES<br>(*n* = 28,517) |  | KEEP (*n* = 127,149) |  |  |  |
| --- | --- | --- | --- | --- | --- | --- | --- |
| Measure | *n* | Weighted |  | Unweighted | Unweighted-GEE | Weighted | Weighted-GEE |
|  |  | % (95% CI) | *n* | % (95% CI) | % (95% CI) | % (95% CI) | % (95% CI) |
| CKD Stage 3 | 2,453 | 6.23<br>(5.82-6.66) | 16,706 | 13.14<br>(12.95-13.33) | 12.83<br>(11.88-13.85) | 8.57<br>(8.30-8.85) | 6.45<br>(5.70-7.28) |
| CKD Stage 3-5 | 2,692 | 6.73<br>(6.30-7.19) | 17,973 | 14.14<br>(13.94-14.33) | 13.86<br>(12.88-14.90) | 9.14<br>(8.86-9.43) | 6.90<br>(6.12-7.78) |
| CKD Stage 4-5 | 239 | 0.51<br>(0.43-0.59) | 1,267 | 1.00<br>(0.94-1.05) | 1.01<br>(0.93-1.09) | 0.55<br>(0.50-0.60) | 0.52<br>(0.42-0.64) |

Age and sex stratified results in Figure 2 show that both age and sex have substantial impacts on CKD endpoints. Both NHANES and KEEP showed low CKD Stage 3 prevalence at younger ages that increased with age for females more than males (KEEP age x sex interaction *P* = 0.037). Stage 4-5 prevalence also increased with age with female versus male differences not uniform over ages (KEEP age x sex interaction *P* = 0.026).

**Figure 2.**
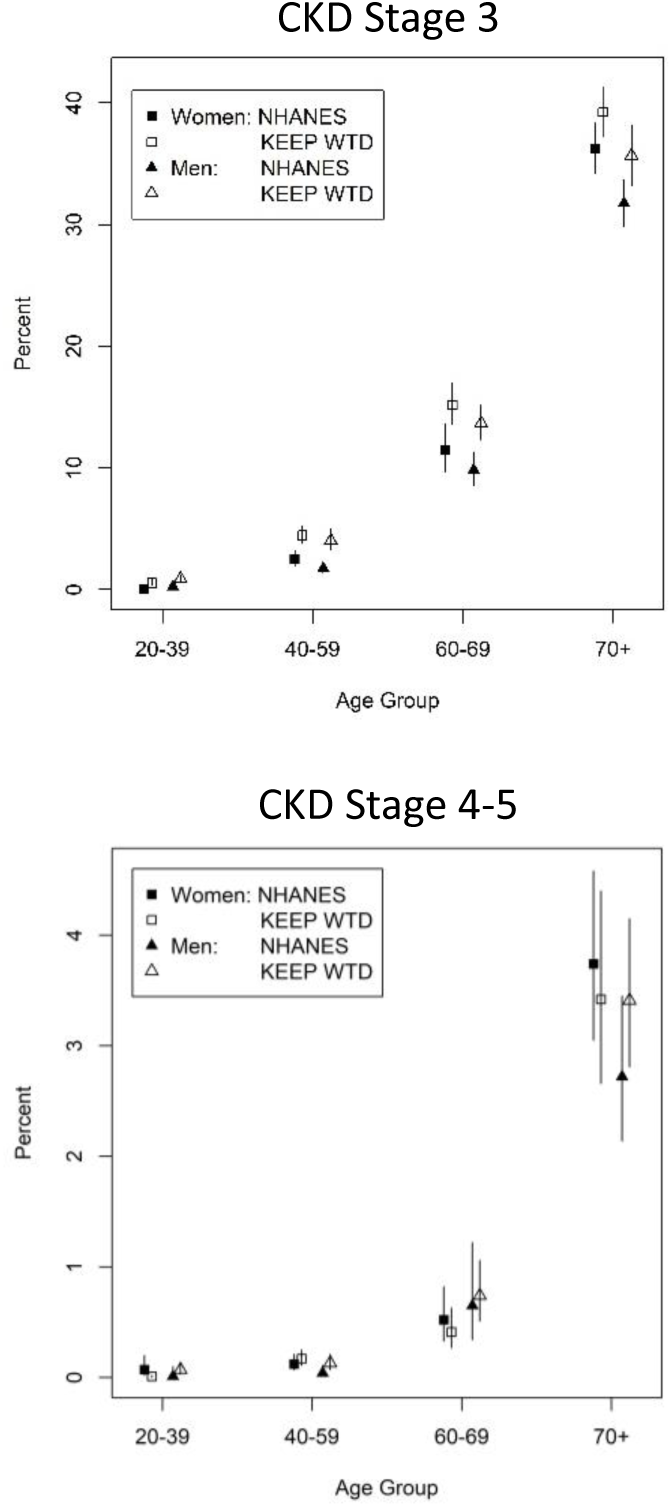
Prevalence of CKD Stage 3 and Stage 4-5 by year screened for KEEP and NHANES. (KEEP - Kidney Early Evaluation Program; NHANES - National Health and Nutrition Examination Survey)

### Temporal Trends in the prevalence of CKD Stage

Prevalence of Stages 3-5 and Stages 4-5 by KEEP year are shown in Figure 1 –3 with and without IPW. NHANES estimates are plotted in the middle of each two-year survey period so that the KEEP estimates falling immediately before and after NHANES estimates are most relevant for comparisons. Tests for non-linear time trends were not significant for CKD Stage 3 (*P* = 0.13), Stages 3-5 (*P* = 0.08) or Stages 4-5 (*P* = 0.07), and no CKD Stage showed a significant positive or negative trend (Supplemental Table 2, all *P*-values > 0.63). Weighted prevalence of CKD Both Stage 3 and Stage 4-5 weighted estimates are close to NHANES in all years with 95% confidence intervals for KEEP trend lines overlapping NHANES point estimates. Weighted KEEP CKD prevalence was slightly higher in early and late years of the series compared to middle years. We used graphical methods to explore whether covariate imbalance in age, sex and race/ethnicity by year may be associated with this extra variation (Supplemental Figures 3 – 5). All three covariates had variation in balance over years, and the pattern for sex most closely matched that seen in weighted KEEP prevalence.

**Figure 3.**
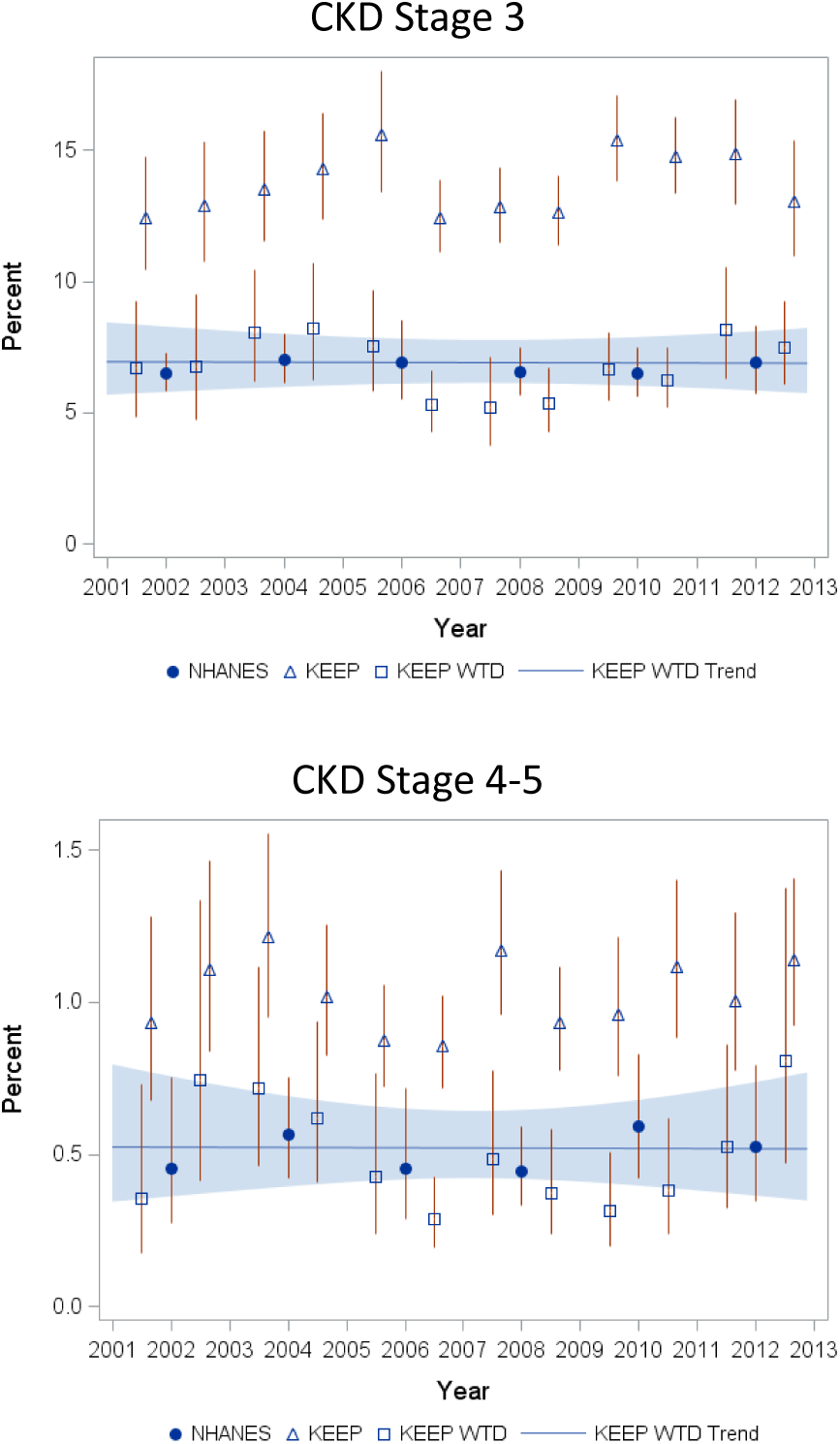
Prevalence (%) of CKD Stage 3 and Stage 4-5 by year screened for KEEP and NHANES. (KEEP - Kidney Early Evaluation Program; NHANES - National Health and Nutrition Examination Survey)

## DISCUSSION

In this paper we explored the use of data from a voluntary, self-selected community disease detection program to model point prevalence and temporal trends for epidemiologic surveillance of CKD in the US over a period of 10 years. By using a national representative survey as a reference, we demonstrate the potential of IPW based adjustments to address the self-selection bias inherent in community based disease detection programs. To our knowledge, this is the first application of this approach in the setting of chronic (kidney) disease and opens up the possibility of using large, readily available samples of convenience in epidemiologic surveillance programs.

The use of self-selected samples for the purpose of tabulating official statistics has been receiving increasing attention in the era of administered surveys distributed over the internet(15,16). Similar to community based disease detection programs, the practical use of these designs suffer from self-selection bias. In recent years, there has been a growing literature regarding the modeling of self-selection effects in such surveys. This literature suggests that a number of approaches, including post-stratification weighting of the observations of participants in the self-selected samples(17) or the direct modeling of the probability of self-selection (“propensity score adjustment”)(13,18,19) against a referent population may allow the valid use of these convenience samples in place of random probability samples. To our knowledge, similar techniques have not been applied to CKD prevalence data. Therefore, we explored both IPW and post-stratification as a novel means to address the self-selection bias in KEEP. Our analyses indicate that even though both methods may substantially decrease this bias, the greater flexibility afforded by the logistic regression model in IPW leads to greater comparability with the sample estimates from NHANES. We postulate that this performance of the IPW is likely to hold true in other health domains outside the field of CKD research. A major strength of our proposed approach to the surveillance of the CKD epidemiology is the use of co-temporal, national, representative cohort of NHANES to calculate the self-selection probabilities in KEEP. The use of such a reference, random probability sample is an integral component of existing approaches for handling selection bias(13,17,20,21). Even if such a reference sample is available though, in order for sample weighting to reduce bias, the covariates used should be strongly correlated with the target population(15) and the mechanism underlying the self-selection process should be one of Missing-At-Random (MAR). In such a case, IPW will allow the unbiased estimation of quantities of interest e.g. point prevalence for impaired eGFR or microalbuminuria (15). In our approach, the rich data collected during KEEP and the in-depth knowledge of the KEEP data collection, the community advertisement and engagement processes led us to consider a-priori plausible subsets of covariates to use for the weighting process, and suggested the need to employ a GEE model to allow for variation among the NKF affiliates where screenings were performed. These factors are likely responsible for the enhanced comparability of the weighted estimates of KEEP to NHANES over the entire period 2001-2012 and at specific points in time during the same time interval.

In terms of practical applications, successfully addressing the bias in KEEP offers the possibility of using this readily available sample of convenience for the purpose of CKD surveillance. In particular, the existing CKD surveillance project about the general US population is based on a limited number of NHANES participants with an eGFR <60 (*n* = 2,700) and eGFR<30 (*n* = 239). Our analyses indicate the potential to expand this dataset almost six-fold by appropriate weighting of the nearly 18,000 KEEP participants with decreased renal function. Notwithstanding the increase in sample size, the ability to combine these datasets affords the opportunity to overcome the pitfalls of each of these two studies when considered in isolation (7,10,22,23). Whereas KEEP suffers from self-selection bias(10,22), NHANES does not represent well the advanced, but also less common stages 3-4 of CKD (7). Such patients are “oversampled” in KEEP so that these studies directly complement each other. Our report illustrates the feasibility of using relatively simple weighting adjustments to make the estimates of the studies directly comparable. Hence, it represents a significant advance over the existing, semi-qualitative use of the two data sources by the nephrology research community. In particular, it was previously stated that the results of KEEP are best understood in “the context of the US population” and in comparison against results from a representative sample (NHANES)(22,24). By applying methodologies to address the self-selection bias in KEEP we took this approach to its logical conclusion and derived a common analytic file that is available for the epidemiologic surveillance of CKD in the general population.

Despite the ability of our methodology to bring about a quantitative agreement between NHANES and weighted versions of KEEP, our analyses have certain limitations. First, recruitment efforts for the KEEP survey was rather heterogeneous over the continental US and may have not reached some population segments. This potential source of bias may not be accounted for in our approach and is a topic under investigation using recruitment information like screening event location, and measures of advertising ‘effort’ as potential covariates. In our analyses we handled these factors indirectly by including the regional affiliates as clusters in the GEE estimation procedure. It is likely that more detailed modeling may have led to more precise estimation of the prevalence of the different stages of CKD. Another limitation concerns the handling of race and ethnicity, which are clinically significant correlates of CKD risk in the source datasets. Whereas NHANES collects detailed race and ethnicity information, the coarseness of classification in public use files likely combines groups with unequal risk. These categories were not entirely congruent with the ones adopted in the KEEP program, thus raising the possibility of residual confounding in our weighting schemes. Finally, adjustment by post-stratification typically relies on a known reference distribution such as from official census statistics.

In summary, we present the first to our knowledge application of self-selection bias correction methodologies for the analysis of data related to the prevalence of CKD in the general US population. We found that two methodologies, i.e. IPW and to a lesser degree post-stratification weighting may be used to render estimates from a self-selected cohort (KEEP) directly comparable to those from a national representative sample (NHANES). Future studies should build on this effort and utilize this novel analytic set to expand the existing national CKD surveillance system. Such efforts may be directed towards understanding the epidemiology of CKD by utilizing the geographic information collected during KEEP so as to build prevalence maps of this challenging, costly chronic disease over both space and time.

## MATERIALS AND METHODS

### Study Populations

We used individual level data from participants in the National Kidney Foundation’s (NKF) Kidney Early Evaluation Program (KEEP) and from participants in the National Health and Nutrition Examination Survey (NHANES). Both KEEP and NHANES are national samples: KEEP is a self-selected sample of adults with elevated risk of kidney disease coordinated by local NKF organizations, and NHANES is a nationally representative sample designed to study health and nutritional status of adults and children in the United States. KEEP used advertising campaigns to attract participants to examinations coordinated by regional affiliates. Advertising targeted participants that were at least 18 years old with risk factors for CKD: high blood pressure, diabetes, or a family history of diabetes or hypertension or kidney disease. We obtained data from NKF for KEEP participants assessed by 48 regional affiliates from 2001 to the end of 2012 (185,511 records, SAS file name keep_saf_2013). Participants were excluded if <20 years of age, on dialysis, were pregnant, had a previous kidney transplant, if eGFR could not be determined, were lacking a valid state of residence or had other invalid demographic variable values. NHANES employs cross-sectional, multi-stage, stratified, cluster probability samples with several subgroups oversampled to improve precision. The subgroups vary by two-year survey period with respect to status race, ethnicity and poverty level. We studied participants attending mobile examination centers (MEC) during six survey periods covering 2001 – 2012.

### Selection Model Variables

Covariates for selection models were chosen if they were related to messages used for KEEP recruitment, if they could be identified in both datasets, and if they received or could be re-encoded similarly in both datasets over the 2001 – 2012 study. Demographic covariates included continuous age, sex, and race/ethnicity (non-Hispanic White, non-Hispanic Black, Mexican/Other Hispanic, Other). Factors related to KEEP recruitment advertising included patient reported diabetes, hypertension, CKD, and family history of CKD, diabetes or hypertension. Other self-reported factors potentially associated with KEEP participation were obesity status, smoking, family history of heart attack, and participant cardiovascular disease (self-report of stroke, congestive heart failure, angina or heart attack). We also included variables for participant socioeconomic status, education and type of insurance.

### CKD Endpoints

Our CKD end points to be estimated from KEEP data after adjusting for self-selection were the single sample (‘point’) prevalence of impaired renal function: CKD Stages 3, eGFR 30 - <60 ml/min/1.73 m^2^, Stages 3-5, eGFR <60 ml/min/1.73 m^2^, and Stage 4-5 CKD, eGFR <30 ml/min/1.73 m^2^. eGFR was estimated using the Chronic Kidney Disease Epidemiology Collaboration (CKD-EPI) equation based on race, sex and serum creatinine(25,26). KEEP serum creatinine values were standardized to the Cleveland Clinic(27) prior to calculation eGFR. Urine creatinine concentrations before 2007 in the NHANES data were standardized to later years using a piece-wise regression adjustment described for NHANES results (28).

### Statistical Analyses

The analyses had two phases: 1) estimation of propensity scores using NHANES and KEEP, and 2) estimation of CKD prevalence. Sampling weights for NHANES were the two-year MEC weights. We conducted analyses to determine whether individual propensity scores for KEEP were sensitive to the approach used in developing the reference population (Supplemental Methods). We created an NHANES summary file composed of weighted reference population frequencies for all cross-tabulations of our selection model covariates. These weighted frequencies considered the complex sampling design and used NHANES PSU, strata, and MEC in calculations. We appended these weighted subgroup frequencies to KEEP data to also estimate propensity scores. Analyses using the two approaches confirmed that propensity scores for KEEP participants were not sensitive how we created a reference population. Using the individual NHANES records and MEC without upscaling to subgroup frequencies for the total reference population avoids the problem of some subgroup cells estimated from very few NHANES observations. Our approach also enables use of continuous variables like age and BMI in estimating propensity scores.

Descriptive summaries of NHANES participants by subgroups (e.g., age, sex, race/ethnicity) were estimated using methods that accounted for the complex sampling design when summarizing the reference population (SURVEYFREQ, SAS v9.4). The population distribution of KEEP for these subgroups were estimated without and with weighting for self-selection.

For propensity score analyses we used boosted the classification and regression tree (CART) methods implemented in the *twang* R-package(29), as this method outperforms logistic regression(30) and can utilize sampling weights. We entered NHANES observations using the MEC weights, and KEEP observations using sampling weights equal to 1.0. The shrinkage parameter was set at 0.001 and number of random trees equaled 20,000. We varied interaction depths from four to six levels and assessed the balance achieved by comparing KEEP weighted frequencies to the expected frequencies based on the NHANES reference population and selected as our propensity score estimation model the one with best agreement between KEEP frequencies expected for the reference population and the weighted frequencies from the boosted CART models. Separate propensity score analyses by two-year NHANES sampling periods estimated weights that tracked changes in KEEP sampling distribution and intensity by regional KEEP affiliates. Achieved covariate balance by NHANES survey period was assessed using unstandardized and standardized differences in proportions(31), and logistic regression was used to assess whether CKD Stage 3-5 prevalence in the unweighted KEEP sample was constant over IPW deciles(32).

We conducted analyses to determine whether individual propensity scores for KEEP were sensitive to the approach used in developing the reference population (Supplemental Methods). We created an NHANES summary file composed of weighted reference population frequencies for all cross-tabulations of our selection model covariates. These weighted frequencies considered the complex sampling design and used NHANES PSU, strata, and MEC in calculations. We appended these weighted subgroup frequencies to KEEP data to also estimate propensity scores. Analyses using the two approaches confirmed that propensity scores for KEEP participants were not sensitive how we created a reference population. Using the individual NHANES records and MEC without upscaling to subgroup frequencies for the total reference population avoids the problem of some subgroup cells estimated from very few NHANES observations. Our approach also enables use of continuous variables like age and BMI in estimating propensity scores.

The second analysis stage had the goal of estimating CKD prevalence by Stage using the KEEP sample but adjusted for self-selection. Unweighted and weighted prevalence and 95% confidence interval are reported (SURVEYFREQ, SAS v9.4) along with population average estimates that account for participants clustered within regional affiliate recruitment and examination programs (GEE, GENMOD, SAS v9.4). Confidence intervals based on robust standard errors are reported. Intercept-only models estimated overall prevalence during 2001-2012 by CKD Stage and during 2003-2012 for albuminuria. Models with fixed effects for age and sex and for KEEP screening year estimated prevalence grouped by these factors. Linear time models and models with restricted cubic splines (*k* = 3 knots) were used to estimate trends in prevalence. Prevalence estimates from NHANES for comparison with weighted KEEP accounted for the complex sampling design but did not use a GEE approach (SURVEYLOGISTIC, SAS v9.4).

### Data availability

NHANES data are available from the survey website at the Center for Disease Control (https://www.cdc.gov/nchs/nhanes/index.htm). The KEEP data that support the findings of this study are available from the National Kidney Foundation (NKF) but restrictions apply to the availability of these data, which were used under license for the current study, and so are not publicly available. Data are however available from the authors upon reasonable request and with permission of NKF.

This secondary analysis of the KEEP and NHANES was approved by the Human Research Protection Office (HRPO) of the University of New Mexico Health Sciences Center (Decision Number HRCC #14-264 on 9/12/2014).

## ACKNOWLEDGMENTS

Support: This work was partially supported by Dialysis Clinic, Inc under the grant #C-3763 Geosurveillance of the Chronic Kidney Disease Epidemic in the US. Preliminary versions of this work were presented in abstract form during Kidney Week 2016 & 2017, Annual Meetings of the American Society of Nephrology and at the GEOMED 2017 meeting held in Porto, Portugal in 2017.

## Author Contributions

All co-authors contributed in this collaborative work: Dr OM programmed the analyses in SAS, created graphs and tables and drafted the Methods and Results of the manuscript. Dr VSP, oversaw statistical analyses and contributed in the Methods section. Dr’s KM, JV, AG contributed their in-depth knowledge of the KEEP and NHANES datasets and contributed sections in Introduction, Methods and Discussion. Dr MU contributed sections in the Introduction and Discussion. Dr CA devised the overall analytic strategy and prepared the Introduction and the Discussion sections of the first draft of the text. All authors, interpreted results and modified the draft of the text over four iterations in order to produce the final version of the manuscript.

## Additional Information

### Competing Interests Statement

The authors declare no competing interests

### Supplementary Information

Supplementary Methods and Results are available from the journal website

